# LRRK2-dependent Rab GTPase phosphorylation in response to endolysosomal damage depends on macrophage differentiation

**DOI:** 10.1101/2020.10.02.323873

**Authors:** Susanne Herbst, Maximiliano G. Gutierrez

## Abstract

The Parkinson’s Disease (PD) kinase LRRK2 is highly expressed in immune cells such as macrophages. In these cells, LRRK2 regulates innate immune pathways and it is activated after membrane damage leading to the phosphorylation of the Rab GTPases Rab8A and Rab10. Due to their wide-range of functions in immunity and tissue remodelling, macrophages *in vivo* are phenotypically heterogeneous. *In vitro* systems are used to differentiate these cells into diverse macrophage subsets to mimic the populations observed *in vivo*. M-CSF and GM-CSF differentiated human blood monocytes are often used to generate monocyte-derived macrophages as a model for tissue macrophages. However, how LRRK2 is activated in different macrophage subsets after membrane damage is unknown.

Here, we report that bone marrow derived macrophages and human monocyte-derived macrophages differentiated with either M-CSF or GM-CSF show different levels of LRRK2 activation after membrane damage. Notably, the membrane damaging agent LLOMe triggered LRRK2-dependent Rab8A and Rab10 phosphorylation primarily in GM-CSF differentiated macrophages. Moreover, LRRK2 and Rab8A were recruited to damaged endolysosomes in GM-CSF differentiated macrophages. Strikingly, GM-CSF differentiated macrophages recruited significantly more CHMP4B and Galectin-3 into damaged endolysosomes. These results suggest that LRRK2-regulated pathways of endolysosomal membrane damage and repair differ between macrophage subsets.

## Introduction

Mutations in Leucine-rich repeat kinase 2 (LRRK2) underlie inherited forms of Parkinson’s Disease (PD) and genome-wide association studies have identified polymorphisms in LRRK2 as a risk factor for Crohn’s Disease and inflammatory forms of leprosy (1). LRRK2 is highly expressed in cells of the immune system, particularly in macrophages, neutrophils and B cells (2). In the last years, research has provided compelling evidence that LRRK2 has an immune function. However, how LRRK2 immune functions relates to PD pathology remains poorly understood. In particular, elevated cytokine levels in the cerebrospinal fluid of PD patients suggest that neuroinflammation constitutes a disease driver (3, 4). As mediators of neuroinflammation, current research predominantly focuses on microglia, the resident macrophage of the brain. However, there are conflicting data regarding the levels of LRRK2 expression in microglia and it has been suggested that abnormal LRRK2 function in peripheral immune cells contributes to neuroinflammation (5). Due to their wide-range of functions in immunity and tissue remodelling, macrophages *in vivo* are phenotypically heterogeneous (6). *In vitro* systems are used to differentiate these cells with different growth factors to mimic the population diversity observed *in vivo*. M-CSF differentiated human blood monocytes are often used to generate monocyte-derived macrophages as a model for tissue-resident macrophages and GM-CSF differentiated monocytes are widely used as a model system for macrophages with tissue remodelling capabilities or alveolar macrophages.

LRRK2 phosphorylates a subset of Rab GTPases which are major regulators of vesicle trafficking (7, 8). As such, LRRK2 has been implicated in a variety of cellular processes, including autophagy, endocytic recycling and lysosomal homeostasis (9–11). LRRK2 regulates lysosomal homeostasis by activating membrane repair mechanisms in response to endomembrane damage and stress (11–13). However, if LRRK2 activation in response to membrane damage is a universal process in macrophage subsets is unknown.

In this report, we investigated if macrophage differentiation impacts LRRK2 activation in response to endomembrane damage. We investigated LRRK2 activation in both human and mouse primary macrophages differentiated with either M-CSF or GM-CSF. Strikingly, GM-CSF-differentiation markedly increased LRRK2 activation in response to endomembrane damage despite similar expression levels of LRRK2 and its substrates Rab8A and Rab10. These findings indicate that LRRK2 function might differ not only between cell types but also within cell type subsets depending on the tissue environment they are exposed, uncovering an avenue of research of LRRK2 regulation and activation.

## Results

To investigate the effect of macrophage differentiation, murine bone marrow was differentiated with either M-CSF or GM-CSF for seven days to generate bone marrow derived macrophages (BMDM). In order to assess LRRK2-dependent Rab GTPase phosphorylation, cells were treated with the endomembrane damaging agent LLOMe and LRRK2 kinase activity was inhibited using the potent inhibitor of LRRK2 kinase activity MLi-2 (14). In GM-CSF-derived murine macrophages, LLOMe treatment resulted in Rab8A and Rab10 phosphorylation (**Fig. 1A**). However, to our surprise, we did not observe Rab GTPase phosphorylation in M-CSF-differentiated macrophages (**Fig. 1A**). We then validated these findings in human CD14 positive blood-derived monocytes macrophages differentiated as above. Similar to what we observed in murine macrophages, LLOMe treatment of GM-CSF differentiated human macrophages resulted in LRRK2-dependent Rab8A and Rab10 phosphorylation (**Fig. 1B-C**). In contrast, Rab GTPase phosphorylation in M-CSF-differentiated macrophages was significantly lower despite similar levels of LRRK2, Rab8A and Rab10 expression (**Fig. 1B-D**). It is of interest to note however, that LRRK2 pS935 phosphorylation was significantly increased in GM-CSF-derived macrophages (Fig. 1B-D).

**Figure 1:**
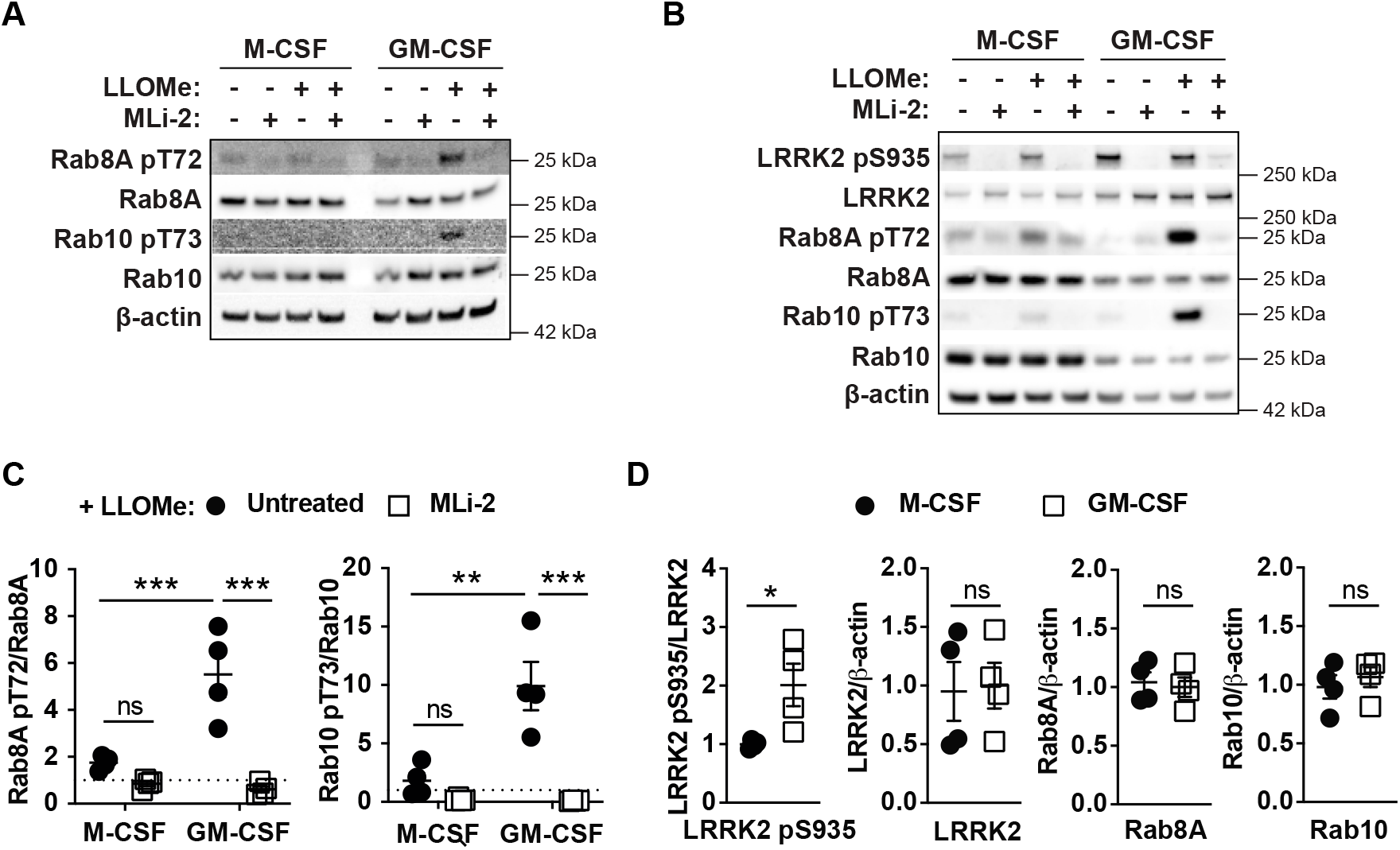
LLOMe-induced LRRK2-dependent Rab8A and Rab10 phosphorylation is primarily detected in GM-CSF differentiated macrophages. **(A)** Murine bone-marrow derived (One out of two independent experiments shown) or **(B)** human blood CD14+ monocytes were differentiated in 50 ng/ml M-CSF or GM-CSF and stimulated with 1 mM LLOMe for 30 min. Where indicated, the cells were pre-treated with 100 nM MLi-2. Cell lysates were probed for LRRK2 pS935, Rab8A pT72 and Rab10 pT73. **(C)** Quantification of Rab phosphorylation in human MDMs, n = 4. All samples were normalised to untreated, non-stimulated cells (indicated by the dotted line). **p<0.01; ***p<0.001 by one-way ANOVA followed by Tukey’s multiple comparisons test. **(D)** LRRK2 pS935 phosphorylation, and LRRK2, Rab8A and Rab10 protein level quantification in human MDMs, n = 4. ns, not significant; \**p*<0.05 by Mann-Whitney test.

LLOMe treatment results in the recruitment of LRRK2 and its substrate Rab8A to damaged endolysosomes (12, 13). We therefore tested LRRK2 and Rab8A positive vesicle formation in response to endolysosomal damage in M-CSF and GM-CSF differentiated BMDM. In agreement with the previous results, we detected LRRK2 recruitment to lysosomes in GM-CSF but not M-CSF differentiated BMDM (**Fig 2A**). Similarly, LRRK2-dependent Rab8A recruitment was significantly more pronounced in GM-CSF differentiated macrophages (**Fig 2B**). Notably, although very rare, some GM-CSF differentiated macrophages already showed significantly higher numbers of Rab8A positive vesicles than M-CSF differentiated macrophages without LLOMe treatment (**Fig. 2B**).

**Figure 2:**
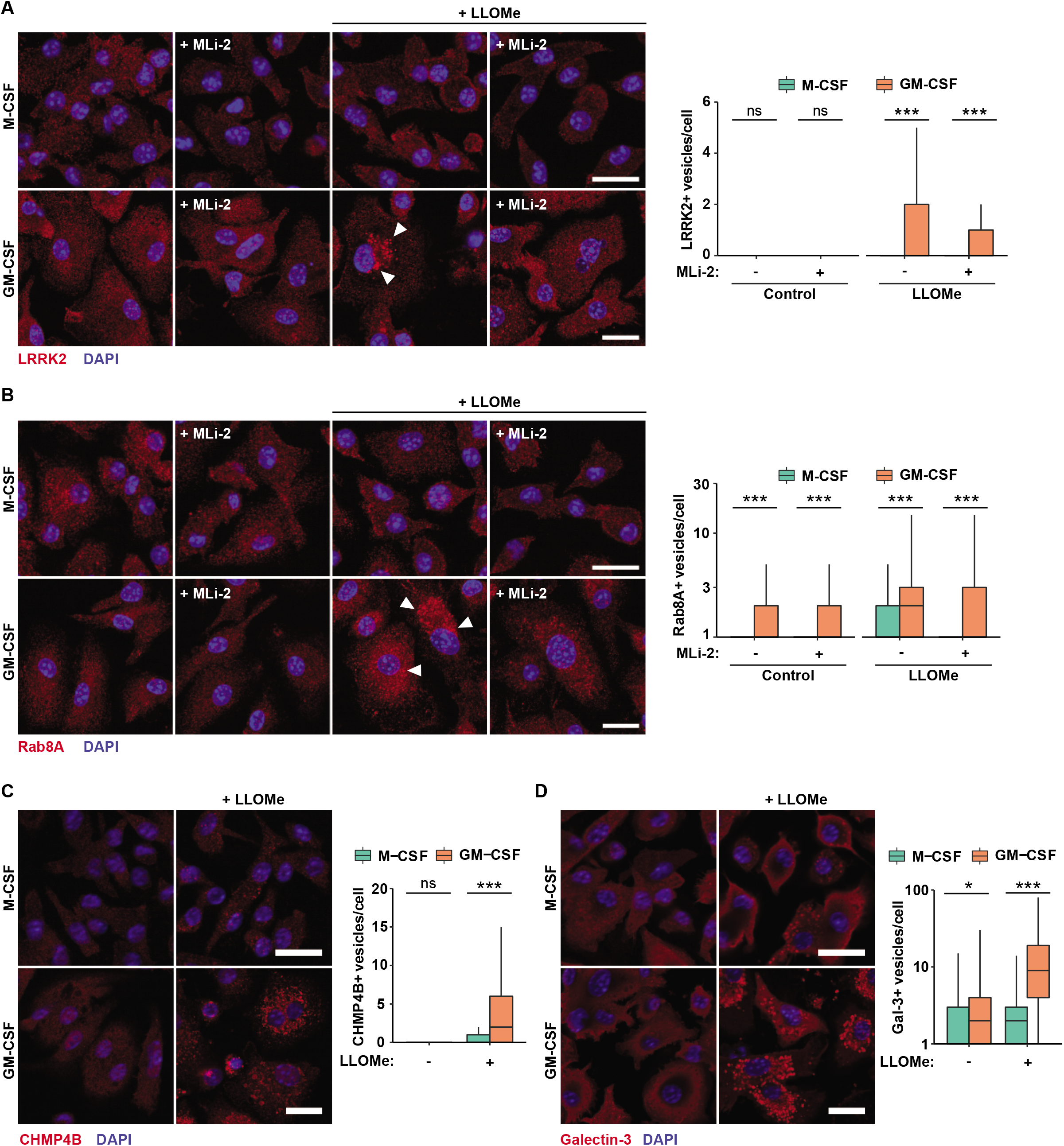
LRRK2 and Rab8A are recruited to damaged endolysosomes in GM-CSF differentiated macrophages. Murine bone-marrow derived cells were differentiated in 50 ng/ml M-CSF or GM-CSF and stimulated with 1 mM LLOMe for 30 min. Where indicated, the cells were pre-treated with 100 nM MLi-2. **(A)** LRRK2, **(B)** Rab8A, (**C**) CHMP4B and (**D**) Galectin-3 positive vesicle formation after endolysosomal damage was assessed by immunofluorescence and high-content imaging. Scale bar = 20 μm. ns, not significant: \**p*<0.05; \*\*\**p*<0.001 by one-way ANOVA followed by Sidak’s multiple comparison’s test.

LRRK2 regulates a membrane repair pathway that involves the ESCRT component CHMP4B and the endomembrane damage sensor Galectin-3 (12). Because the recruitment of LRRK2 and Rab8A to damaged endolysosomes was prominently present in GM-CSF differentiated macrophages, we tested if this was due to alterations in endomembrane repair. For that, we analysed both CHMP4B and Galectin-3 recruitment to damaged endolysosomes in M-CSF and GM-CSF differentiated macrophages. LLOMe treatment resulted in the formation of CHMP4B and Galectin-3 positive vesicles in both M-CSF and GM-CSF differentiated macrophages (**Fig. 2C-D**). However, in agreement with a role for LRRK2 in ESCRT and Galectin-3 function, GM-CSF differentiated macrophages recruited significantly more CHMP4B and Galectin-3 after endomembrane damage (**Fig. 2C-D)**. In the same way as we detected Rab8A positive vesicles in the absence of LLOMe treatment, we also detected increased numbers of Galectin-3 positive vesicles in GM-CSF differentiated control macrophages (**Fig. 2D**). Taken together, these findings highlight that macrophage differentiation impacts LRRK2 activation and endomembrane repair in response to endolysosomal damage.

## Discussion

Macrophages have taken centre stage in the quest to identify immune functions of LRRK2, particularly because resident macrophages in form of microglia can be found in the brain. We have recently shown that endomembrane damage in macrophages constitutes a trigger for LRRK2 activation resulting in Rab GTPase phosphorylation and ESCRT-mediated membrane repair (12). Here, we demonstrate that macrophage differentiation modulates the ability of macrophages to activate LRRK2 and the kind of response that is mounted to endomembrane damage. We did not detect pronounced differences in LRRK2 protein levels in the two macrophage subsets, which suggests that LRRK2 might execute different functions, dependent on the cell type it is expressed in or that LRRK2 activity can be regulated by extracellular cues. Interestingly, Xu *et al.* reported no apparent differences between M-CSF and GM-CSF differentiated human or mouse macrophages after 48 h of LPS treatment and that IFN-γ up-regulated LRRK2 protein levels in both M-CSF and GM-CSF differentiated macrophages increasing basal levels of Rab10 phosphorylation (15). This indicates that extrinsic factors such as cytokines or pathogen-associated molecular patterns like LPS determine if LRRK2 is activated in response to endomembrane damage. This leads to the question which signalling pathways regulate LRRK2 activation and if there are post-transcriptional modification on LRRK2 that determine activation. LRRK2 pS935 is a known Toll-like receptor responsive phosphorylation side (16) which has been shown to regulate LRRK2 localisation without directly affecting LRRK2 kinase activity (17). We observed increased LRRK2 pS935 phosphorylation in GM-CSF differentiated macrophages, which requires further investigation to understand its functional implications.

There is considerable interest in using the phosphorylation of Rab GTPases as a read-out to monitor LRRK2 activation in PD patients. Our study suggests that the development of Rab GTPase phosphorylation as a biomarker of LRRK2 activity needs to be carefully analysed and that potential differentiation and activation protocols need to be standardised.

Importantly, data shown here highlights that mechanisms of endomembrane damage and repair are likely different in different macrophage subsets. We hypothesize that *in vivo*, different populations of macrophages and likely other myelocytic cells will trigger different inflammatory outcomes due to these differences in the ability to repair membrane damage after infection or protein aggregation. GM-CSF is also used to differentiated monocytes into monocyte-derived dendritic cells (18), a cell type that has to balance lysosomal degradation with preservation of antigens for presentation to T cells (19). Regulating lysosomal permeability could be one way of achieving this balance (20). If LRRK2 contributes to dendritic cell function or any other immune cell such as B cells and neutrophils remains to be elucidated and could significantly contribute to our understanding of LRRK2-dependent PD pathology.

## Experimental procedures

### Human monocyte-derived macrophage isolation and differentiation

Monocytes were isolated from leukocytes cones supplied by the NHS blood and transplant service. First, red blood cells were removed by centrifugation on Ficoll-Paque (28-4039-56 AD; GE Healthcare) and recovered PBMCs were washed twice in PBS-0.5 mM EDTA to remove platelets. Monocytes were isolated from PBMCs using a magnetic cell separation system with anti-CD14 mAb-coated microbeads (130-050-201; Miltenyi Biotec). CD14-positive monocytes were differentiated in complete RPMI 1640 medium (Gibco) supplemented with 10 % heat-inactivated FCS and 50 ng/ml GM-CSF (130-093-862; Miltenyi Biotec). Cells were cultured at 37°C under a humidified 5% CO2 atmosphere for 6 days and the medium was replaced at day 3. On day 6, the cells were washed and detached with 0.5 mM EDTA in ice-cold PBS and plated in RPMI 1640 medium containing 10 % heat-inactivated FCS.

### Murine bone marrow-derived macrophage differentiation

Bone marrow (BM) was isolated from the hind legs of 8 −10 weeks old mice. Red cell lysis was performed in red blood cell lysing buffer (Hybri-Max, Sigma) for 10 min at RT, before plating the cells in petri dishes in RPMI 1640 medium containing 10 % heat-inactivated FCS and 50 ng/ml M-CSF (BioLegend) or 50 ng/ml GM-CSF (BioLegend). Cells were cultured at 37°C under a humidified 5 % CO2 atmosphere for 6 days and the medium was replaced at day 3. On day 6, the cells were washed and detached with 0.5 mM EDTA in ice-cold PBS and plated in RPMI 1640 medium containing 10 % heat-inactivated FCS.

### LLOMe treatment and LRRK2 kinase inhibition

A 333 mM stock of LLOMe (Cat# 4000725, Bachem) was prepared in ethanol and frozen at −20°C in tightly sealed tubes. For LLOMe treatment, the medium was replaced with RPMI 1640 medium containing 10 % heat-inactivated FCS and 1 mM of LLOMe and the cells were stimulated for 30 min. 0.3 % ethanol in RPMI 1640 medium containing 10 % heat-inactivated FCS was used in all control samples. Macrophages were treated with MLi-2 (Cat# 5756/10, Tocris) at 0.1 μM for 1 h before treatment with LLOMe where indicated. MLi-2 was present during LLOMe treatment.

### Antibodies for Immunofluorescence and Western Blot

Antibodies used in this study were anti-Rab8A (6975), anti-Rab10 (8127) and anti-β-actin-HRP (12262) from Cell Signaling; anti-Rab8A pT72 (ab230260), anti-Rab10 pT73 (ab230261), anti-LRRK2 pS935 (ab133450), and anti-LRRK2 for immunofluorescence (ab133474) from Abcam; anti-LRRK2 for Western Blot (N241A/34) from NeuroMab; anti-Galectin-3-AF647 (125408) from BioLegend and anti-CHMP4B (13683-1-AP) from ProteinTech.

### Western Blotting

For lysis, cells were washed once with ice-cold PBS, harvested in PBS and lysed 1x RIPA lysis buffer (Milipore) containing a protease and phosphatase inhibitor cocktail (ThermoFisher Scientific). The samples were boiled at 70°C for 10 min in LDS sample buffer and reducing agent (NuPAGE, Life Technologies) and run on a NuPAGE 4-12% Bis-Tris gel (Life Technologies). The gels were transferred onto a PVDF membrane by wet transfer. The membranes were blocked in 5 % semi-skinned milk in TBS-T (TBS, 0.1 % Tween-20). The membranes were incubated with primary antibodies in 5 % semiskinned milk in TBS-T at 4 °C overnight and with the secondary antibodies in 5 % skimmed milk in TBS-T for 1 hr at room temperature. Western blots were quantified by densitometry using ImageJ.

### Indirect immunofluorescence

Cells seeded in 96-well plates (Cell Carrier Ultra, Perkin Elmer) were fixed with 4 % methanol-free PFA (15710, Electron Microscopy Sciences) in PBS for 15 min at 4 °C. The samples were quenched with 50 mM NH4Cl (A9434, Sigma-Aldrich) in PBS for 10 min at room temperature and permeabilized with 0.3 % TritonX-100, 5 % FCS in PBS for 20 min. For LRRK2 and CHMP4B immunofluorescence, cells were permeabilized in ice-cold methanol for 10 min at −20 °C, followed by one wash with PBS and blocking in 5 % FCS in PBS for 20 min at RT.

Primary antibodies were diluted in PBS containing 5 % FCS and incubated for 1 hr at RT. The samples were washed 3 times in PBS and when required, the secondary antibody was added in the same way as the primary antibody (anti-rabbit-Alexa fluor 488, Alexa fluor 568 or Alexa fluor 633, Invitrogen) for 45 min at room temperature. After 3 more washes with PBS, nuclear staining was performed using 300 nM DAPI (Life Technologies, D3571) in PBS for 10 min. Images were acquired in PBS on an Opera Phenix high-content screening system in confocal mode using the 63x objective, 2x binning (PerkinElmer).

### Image analysis

Images acquired using the Opera Phenix high-content screening system were analysed using Harmony analysis software 4.9 (PerkinElmer). First, cells were segmented using the DAPI nuclear signal. Vesicles were identified using the “Spots” building block (global maximum), resulting in a “Number of Spots per cell” read-out parameter. For each experiment, at least 2000 cells were analysed per treatment condition.

## Data availability

The authors declare that this study did not produce any primary datasets.

## Acknowledgements

We would like to thanks all members of the Host-Pathogen Interactions in Tuberculosis laboratory for insightful discussions.

## Author contributions

SH and MGG conceived the project. SH performed all the experiments and analysed the data. SH wrote the paper and prepared the figures with input from MGG.

## References

1. Berwick, D. C., Heaton, G. R., Azeggagh, S., and Harvey, K. (2019) LRRK2 Biology from structure to dysfunction: Research progresses, but the themes remain the same. Mol. Neurodegener. 14, 1–22

2. Herbst, S., and Gutierrez, M. G. (2019) LRRK2 in Infection: Friend or Foe? ACS Infect. Dis. 5, 809–815

3. Schröder, J. B., Pawlowski, M., Meyer zu Hörste, G., Gross, C. C., Wiendl, H., Meuth, S. G., Ruck, T., and Warnecke, T. (2018) Immune Cell Activation in the Cerebrospinal Fluid of Patients With Parkinson’s Disease. Front. Neurol. 9, 1081

4. Chen, X., Hu, Y., Cao, Z., Liu, Q., and Cheng, Y. (2018) Cerebrospinal Fluid Inflammatory Cytokine Aberrations in Alzheimer’s Disease, Parkinson’s Disease and Amyotrophic Lateral Sclerosis: A Systematic Review and Meta-Analysis. Front. Immunol. 9, 2122

5. Wallings, R. L., Herrick, M. K., and Tansey, M. G. (2020) LRRK2 at the Interface Between Peripheral and Central Immune Function in Parkinson’s. Front. Neurosci. 14, 443

6. Gordon, S., and Plüddemann, A. (2017) Tissue macrophages: heterogeneity and functions. BMC Biol. 15, 53

7. Martin Steger, Francesca Tonelli, Genta Ito, Paul Davies, Matthias Trost, Melanie Vetter, Stefanie Wachter, Esben Lorentzen, Graham Duddy, Stephen Wilson, Marco AS Baptista, Brian K Fiske, Matthew J Fell, John A Morrow, Alastair D Reith, Dario R Alessi, and Matthias Mann (2016) Phosphoproteomics reveals that Parkinson’s disease kinase LRRK2 regulates a subset of Rab GTPases. Elife. 10.7554/eLife.12813

8. Steger, M., Diez, F., Dhekne, H. S., Lis, P., Nirujogi, R. S., Karayel, O., Tonelli, F., Martinez, T. N., Lorentzen, E., Pfeffer, S. R., Alessi, D. R., and Mann, M. (2017) Systematic proteomic analysis of LRRK2-mediated Rab GTPase phosphorylation establishes a connection to ciliogenesis. Elife. 6, e31012

9. Manzoni, C., Mamais, A., Dihanich, S., Abeti, R., Soutar, M. P. M., Plun-Favreau, H., Giunti, P., Tooze, S. a., Bandopadhyay, R., and Lewis, P. a. (2013) Inhibition of LRRK2 kinase activity stimulates macroautophagy. Biochim. Biophys. Acta - Mol. Cell Res. 1833, 2900–2910

10. Gómez-Suaga, P., Rivero-Ríos, P., Fdez, E., Blanca Ramírez, M., Ferrer, I., Aiastui, A., López De Munain, A., and Hilfiker, S. (2014) LRRK2 delays degradative receptor trafficking by impeding late endosomal budding through decreasing Rab7 activity. Hum. Mol. Genet. 23, 6779–6796

11. Eguchi, T., Kuwahara, T., Sakurai, M., Komori, T., Fujimoto, T., Ito, G., Yoshimura, S.-I., Harada, A., Fukuda, M., Koike, M., and Iwatsubo, T. (2018) LRRK2 and its substrate Rab GTPases are sequentially targeted onto stressed lysosomes and maintain their homeostasis. Proc. Natl. Acad. Sci. 115, E9115–E9124

12. Herbst, S., Campbell, P., Harvey, J., Bernard, E. M., Papayannopoulos, V., Wood, N. W., Morris, H. R., and Gutierrez, M. G. (2020) LRRK2 activation controls the repair of damaged endomembranes in macrophages. EMBO J. 10.15252/embj.2020104494

13. Bonet-Ponce, L., Beilina, A., Williamson, C. D., Lindberg, E., Kluss, J. H., Saez-Atienzar, S., Landeck, N., Kumaran, R., Mamais, A., Bleck, C. K. E., Li, Y., and Cookson, M. R. (2020) LRRK2 mediates tubulation and vesicle sorting from membrane damaged lysosomes. bioRxiv. 10.1101/2020.01.23.917252

14. Fell, M. J., Mirescu, C., Basu, K., Cheewatrakoolpong, B., DeMong, D. E., Ellis, J. M., Hyde, L. A., Lin, Y., Markgraf, C. G., Mei, H., Miller, M., Poulet, F. M., Scott, J. D., Smith, M. D., Yin, Z., Zhou, X., Parker, E. M., Kennedy, M. E., and Morrow, J. A. (2015) MLi-2, a Potent, Selective, and Centrally Active Compound for Exploring the Therapeutic Potential and Safety of LRRK2 Kinase Inhibition. J. Pharmacol. Exp. Ther. 355, 397–409

15. Xu, E., Boddu, R., Abdelmotilib, H. A., Kelly, K., Sokratian, A., Harms, A. S., Schonhoff, A. M., Bryant, N., Harmsen, I. E., Schlossmacher, M. G., Chandra, S., Krendelshchikova, V., Liu, Z., and West, A. B. (2020) Pathologic α-Synuclein Species Activate LRRK2 in Pro-Inflammatory Monocyte and Macrophage Responses. bioRxiv. 10.1101/2020.05.04.077065

16. Dzamko, N., Inesta-Vaquera, F., Zhang, J., Xie, C., Cai, H., Arthur, S., Tan, L., Choi, H., Gray, N., Cohen, P., Pedrioli, P., Clark, K., and Alessi, D. R. (2012) The IkappaB Kinase Family Phosphorylates the Parkinson’s Disease Kinase LRRK2 at Ser935 and Ser910 during Toll-Like Receptor Signaling. PLoS One. 7, e39132

17. Marchand, A., Drouyer, M., Sarchione, A., Chartier-Harlin, M.-C., and Taymans, J.-M. (2020) LRRK2 Phosphorylation, More Than an Epiphenomenon. Front. Neurosci. 14, 527

18. Helft, J., Böttcher, J., Chakravarty, P., Zelenay, S., Huotari, J., Schraml, B. U., Goubau, D., and Reis e Sousa, C. (2015) GM-CSF Mouse Bone Marrow Cultures Comprise a Heterogeneous Population of CD11c+MHCII+ Macrophages and Dendritic Cells. Immunity. 42, 1197–1211

19. Savina, A., and Amigorena, S. (2007) Phagocytosis and antigen presentation in dendritic cells. Immunol. Rev. 219, 143–156

20. Kozik, P., Gros, M., Itzhak, D. N., Joannas, L., Heurtebise-Chrétien, S., Krawczyk, P. A., Rodríguez-Silvestre, P., Alloatti, A., Magalhaes, J. G., Del Nery, E., Borner, G. H. H., and Amigorena, S. (2020) Small Molecule Enhancers of Endosome-to-Cytosol Import Augment Anti-tumor Immunity. Cell Rep. 32, 107905

